# Modulation of the vesicle code transmitting the visual signal in the retina

**DOI:** 10.1101/2020.04.22.056119

**Authors:** José Moya-Díaz, Ben James, Leon Lagnado

## Abstract

Multivesicular release (MVR) allows retinal bipolar cells to transmit visual signals as changes in both the rate and amplitude of synaptic events. How do neuromodulators reguate this vesicle code? By imaging larval zebrafish, we find that the variability of calcium influx is a major source of synaptic noise. Dopamine increases synaptic gain up to 15-fold while Substance P reduces it 7-fold, both by acting on the presynaptic calcium transient to alter the distribution of amplitudes of multivesicular events. An increase in gain is accompanied by a decrease in the temporal precision of transmission and a reduction in the efficiency with which vesicles transfer visual information. The decrease in gain caused by Substance P was also associated with a shift in temporal filtering from band-pass to low-pass. This study demonstrates how neuromodulators act on the synaptic transformation of the visual signal to alter the way information is coded with vesicles.

## Introduction

Visual processing in the retina begins when photoreceptors convert changes in light intensity into graded changes in membrane potential. These analogue signals are subsequently transformed through a number of different microcircuits before conversion into the digital form of spikes that retinal ganglion cells (RGCs) transmit to higher centres (Masland, 2012). The only bridge between the photoreceptors and RGCs are bipolar cells which transmit signals through ribbon-type synapses specialized for sensory communication (Asari and Meister, 2012; Lagnado and Schmitz, 2015). Because all visual information flows through these synapses, the presynaptic compartment of bipolar cells provides a key control point for a variety of retinal computations, including adaptation to contrast (Demb, 2008; Nikolaev et al., 2013), temporal filtering (Baccus, 2007; Baden et al., 2014; Suh and Baccus, 2014), generation of centre-surround antagonism (Buldyrev and Taylor, 2013) and detection of orientation (Johnston et al., 2019). Here we ask how neuromodulators act on this computational unit to alter the way synaptic vesicles are used to encode the visual signal.

Using live zebrafish expressing the glutamate reporter iGluSnFR (Marvin et al., 2013), it has recently become possible to resolve single glutamatergic vesicles as they are released from individual active zones (James et al., 2019). A fundamental insight provided by this optical approach is that the ribbon synapses of bipolar cells discretize the analogue signal arriving down the axon into ten or more output values through a process of multivesicular release (MVR), in which two or more vesicles release their contents synchronously (Singer et al., 2004). As a consequence, bipolar cells do not transmit information through a binary code involving two symbols (zero or one vesicle), but instead use multiple symbols of different amplitude with information being represented as changes in both the rate and distribution (James et al., 2019). MVR may also be an aspect of the vesicle code operating in other parts of the brain, being a feature of ribbon synapses in auditory hair cells (Goutman and Glowatzki, 2007; Li et al., 2014) as well as conventional synapses in the hippocampus, cortex and cerebellum (Conti and Lisman, 2003; Huang et al., 2010; Molnar and Werblin, 2007; Rudolph et al., 2011; Vaden et al., 2019).

The demonstration that MVR encodes visual information requires us to re-examine how the synaptic compartment of bipolar cells recodes the visual signal for chemical transmission. MVR is likely to be regulated both by the intrinsic electrical properties of the terminal and by extrinsic signals generated by the circuitry of the inner retina. Although bipolar cells have traditionally been considered passive neurons, it is now clear that the voltage-dependent calcium channels controlling transmitter release can also transform the visual signal by generating damped oscillations (Burrone and Lagnado, 1997) or calcium spikes of low amplitude (Dreosti et al., 2011). These electrical events are in turn modulated by fast inhibition, both GABAergic and glycinergic, received from amacrine cells that synapse directly onto the presynaptic compartment (Diamond, 2017). But amacrine cells also regulate signal flow on longer time-scales, by releasing neuromodulators such as dopamine, somatostatin and substance P, which can either increase or decrease the gain of synaptic transmission (Yazulla and Studholme, 2001). For instance, electrophysiological experiments in isolated bipolar cells demonstrate that dopamine potentiates L-type calcium channels within the synaptic compartment (Esposti et al., 2013; Heidelberger and Matthews, 1994), while Substance P inhibits them (Ayoub and Matthews, 1992). How such neuromodulators act on the intact retinal circuit is less clear. To what degree do they regulate synaptic strength? And do they alter the way synaptic vesicles are used to transmit a visual signal?

To investigate how retinal neuromodulators alter transmission of the visual signal we used zebrafish to image both glutamate release and the presynaptic calcium signal in bipolar cells. Dopamine increases the contrast sensitivity of individual active zones by increasing both the frequency and average amplitude of release events, and this occurs through two presynaptic mechanisms: an increase in the average size of the calcium transient and an increase in the efficiency with which calcium triggers vesicle release. Substance P acts in the opposite direction, decreasing contrast sensitivity by reducing both the frequency and amplitude of release events, but this decrease in synaptic gain is accompanied by an increase in the temporal precision of multivesicular events and an increase in the efficiency with which vesicles are used to transfer information about stimulus contrast. We also identify variability in presynaptic calcium signals as a major source of noise in transmission of visual information. Together, these results reveal how retinal neuromodulators act on the process of multivesicular release to alter the strength, temporal precision and efficiency of synapses transmitting the visual signal.

## Results

### Variability in presynaptic calcium signals contributes to variability in vesicle output

Sensory circuits are “noisy” in that they do not generate a stereotyped response to a given stimulus {Faisal, 2008 #154}. The process of synaptic transmission is an important contributor to such noise and in the retina determines the detectability of dim signals {Field, 2002 #155; Grimes, 2014 #133}. Recent work on the ribbon synapses of bipolar cells has demonstrated that the process of multivesicular release is an important aspect of synaptic noise in the retina (James et al., 2019) but the underlying mechanisms are not clear. Does this variable amplitude of multivesicular events simply reflect the stochastic properties of the processes triggering vesicle fusion (Fatt and Katz, 1952; Yang and Xu-Friedman, 2013)? Or do the electrical events that determine calcium influx also vary from trial-to-trial? To investigate this question, we began by comparing glutamate release measured with iGluSnFR (Fig. 1A) and the presynaptic calcium signal measured with SyGCaMP6f (Fig. 1B).

Examples of glutamate transients at an individual active zone are shown in Fig. 1C, elicited using a full-field stimulus modulated at 5 Hz. The average size of the uniquantal event was measured as the first peak in the distribution of event amplitudes (Fig. S1 and S2), from which we made a maximum likelihood estimate of the number of quanta comprising each event (Q_e_), as described in Methods. In a sample of 37 synapses (13 ON and 24 OFF), a stimulus contrast of 20% failed to elicit an exocytic event in 75 ± 4% of stimulus cycle (Fig. 1C) but when an event did occur, the average number of quanta it contained (Q_e_) was 1.7 ± 0.15. Increasing the contrast to 100% increased both the frequency of synaptic events (failure rate of 16 ± 2%) and Q_e_ (3.4 ± 0.25). The distribution of amplitudes at 20%, 60% and 100% contrast are compared in Fig. 1D, from which it can be seen that all three overlapped significantly. A change in contrast was therefore represented, at least in part, by a change in the distribution of amplitudes of multivesicular events.

**Figure 1.**
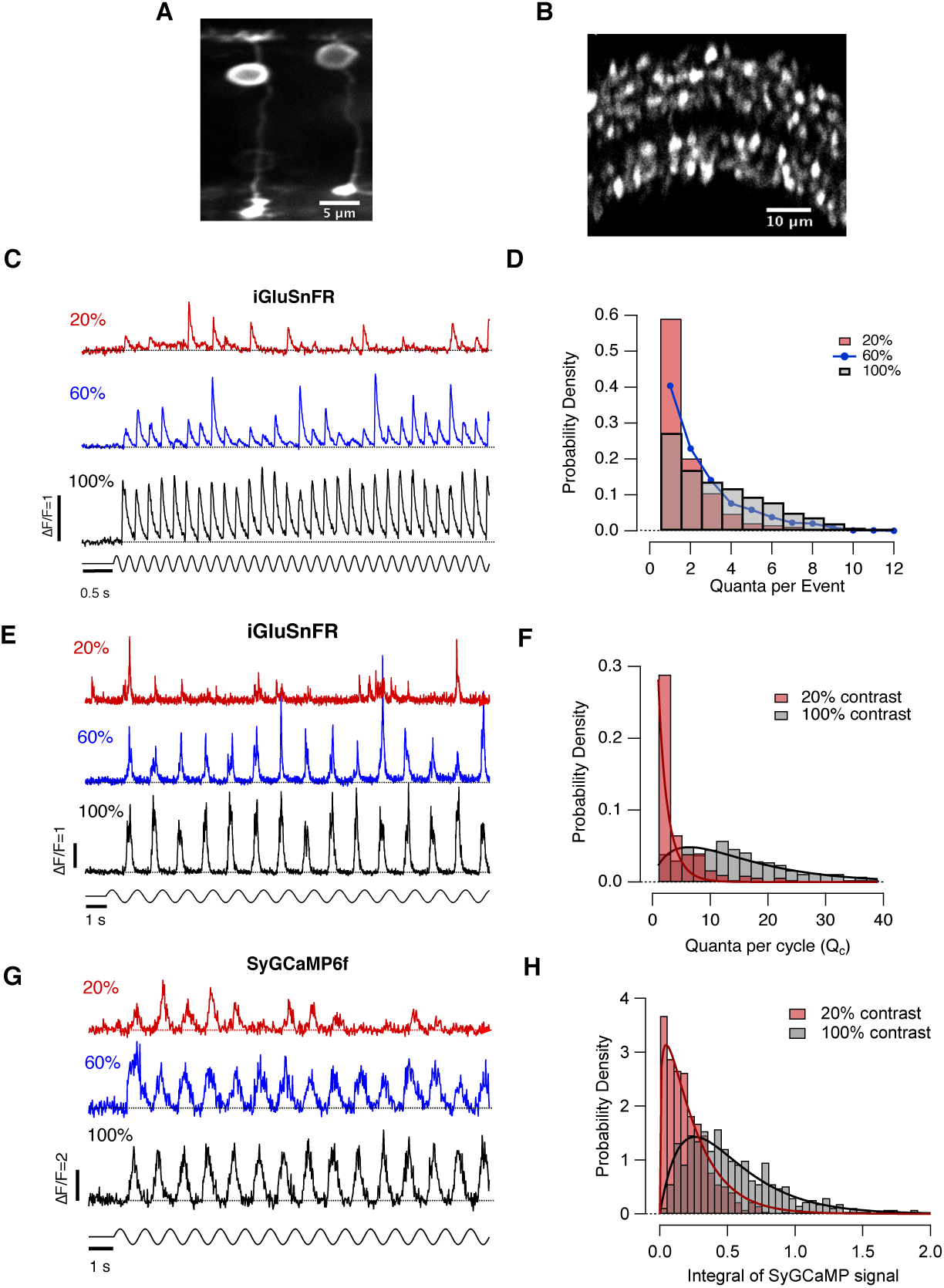
Variability in presynaptic calcium signals and multivesicular release. **A.** Multiphoton section through the eye of a zebrafish larva (7 dpf) expressing the glutamate reporter iGluSnFR in a subset of bipolar cells. No other terminals were in the vicinity. **B.** Expression of the synaptic calcium reporter SyGCaMP6f in synaptic terminals of all bipolar cells through the inner plexiform layer. **C.** Glutamate transients at an individual active zone triggered by stimulus contrasts of 20%, 60% and 100% (full-field sine wave at 5 Hz). Note the variable amplitude and the large number of stimulus cycles failing to elicit a response at 20% contrast. **D.** Changes in the distribution of the number of quanta per event (Q_e_) elicited at 20%, 60% and 100% contrast (n = 37 synapses). Higher contrasts shift the distribution toward larger multiquantal events. **E.** Glutamate transients at a stimulus frequency of 1 Hz. At higher contrasts, large bursts of vesicles are released. **F.** Changes in the distribution of the number of quanta per cycle (Q_C_) elicited at 20% and 100% contrast (n = 10 synapses). The smooth lines are best-fits of the probability density function of the gamma distribution

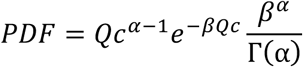

where *α* and *β* are rate and shape parameters, respectively, and Γ(α) is the gamma function. At 20% contrast, *α* = 0.71 and *β* = 0.45. At 100% contrast, *α* = 1.74 and *β* = 0.12. **G.** Synaptic calcium transients at a single terminal triggered by stimulus contrasts of 20%, 60% and 100% (full-field sine wave at 1 Hz). Note the variability in the size of calcium transients and the high frequency of failures at 20% contrast. **H.** Changes in the distribution of calcium transients elicited at 20% and 100% contrast (n = 450 synapses). The smooth lines are best-fits of the probability density function of the gamma distribution. At 20% contrast, *α* = 1.23 and *β* = 5.08. At 100% contrast, *α* = 2.02 and *β* = 3.90.

Variations in the amplitude of synaptic events were quantified as the Coefficient of Variance (CV), the ratio of the standard deviation of Q_e_ to the mean. At conventional synapses, CV has been measured electrophysiologically using the peak amplitude of post-synaptic currents, with mean values of 0.5 at hippocampal synapses (Bekkers et al., 1990) and 0.23 at connections between mossy fibers and granule cells (Silver et al., 1996). One source of such variability in quantal amplitude is thought to be variations in the size of synaptic vesicles (Bekkers et al., 1990) so it is not surprising that MVR in bipolar cells made the variability of synaptic events significantly larger, with CV ranging between 0.77 ± 0.07 at 20% contrast and 2.70 ± 0.26 at 100%.

Although a 5 Hz stimulus allowed exocytic events to be distinguished from each other, it did not allow a comparison with calcium signals measured with SyGCaMP6f because the relatively slow dynamics of the calcium reporter did not allow the signal to recover between cycles. We therefore lowered the stimulus frequency to 1 Hz, which allowed direct comparison of the *total* number of vesicles released during each cycle (Q_c_; Fig. 1E; Supplementary Figure 2) and the integral of the calcium transient (Ca_c_; Fig. 1G). The average value of Q_c_ was 3.8 ± 1.8 vesicles at 20% contrast, increasing to 13.1 ± 2.9 vesicles at 100%. The distribution of these events is shown in Fig. 1F. Surprisingly, the calcium signal driving vesicle fusion was also highly variable: ∼16% of cycles failed to elicit a detectable transient at 20% contrast and Ca_c_ displayed a CV of 0.93 ± 0.05 (red trace in Fig. 1G; histogram in Fig. 1H). The distribution of Ca_c_ measured at 20% and 100% contrast overlapped significantly, in a manner qualitatively similar to the distributions of Q_c_. These results reveal that the electrical events determining calcium influx vary significantly from trial to trial, especially when driven by a stimulus of low contrast, providing a possible explanation for the wide dispersion in the amplitude of MVR events.

### Endogenous dopamine potentiates MVR by increasing the size of calcium transients

Might variations in the activation of calcium channels contribute to variability in synaptic output? The resting potential of bipolar cells sits very close to the threshold for activation of L-type calcium channels, causing regenerative “calcium spikes” to be sensitive to small voltage fluctuations (Baden et al., 2011; Burrone and Lagnado, 1997). We began testing the role of these regenerative mechanisms by manipulating dopamine signaling because this neuromodulator has been shown to lower the voltage threshold for activation of calcium channels (Esposti et al., 2013; Heidelberger and Matthews, 1994).

Drugs activating or inhibiting D1 dopamine receptors were similarly effective in 23/27 OFF and 13/15 ON synapses, and results from the two types are therefore collected together in Fig. 2 (see also Supplementary Fig. 3). Injection of the D1 antagonist SCH 23390 directly into the eye (estimated final concentration of 1 μM) strongly suppressed both presynaptic calcium transients and glutamate release elicited by stimuli of 20% and 60% contrast (Fig. 2A-D). Activation of dopamine signaling above normal levels by injection of the D1 agonist ADTN (estimated concentration of 0.2 μM) had opposite effects, increasing both calcium influx and Q_c_, the total number of vesicles released per cycle of the stimulus. These results demonstrate that endogenous dopamine acted through D1 receptors to strongly potentiate transmission of the visual signal and this occurred, at least in part, by augmenting calcium influx into the presynaptic compartment of bipolar cells.

**Figure 2.**
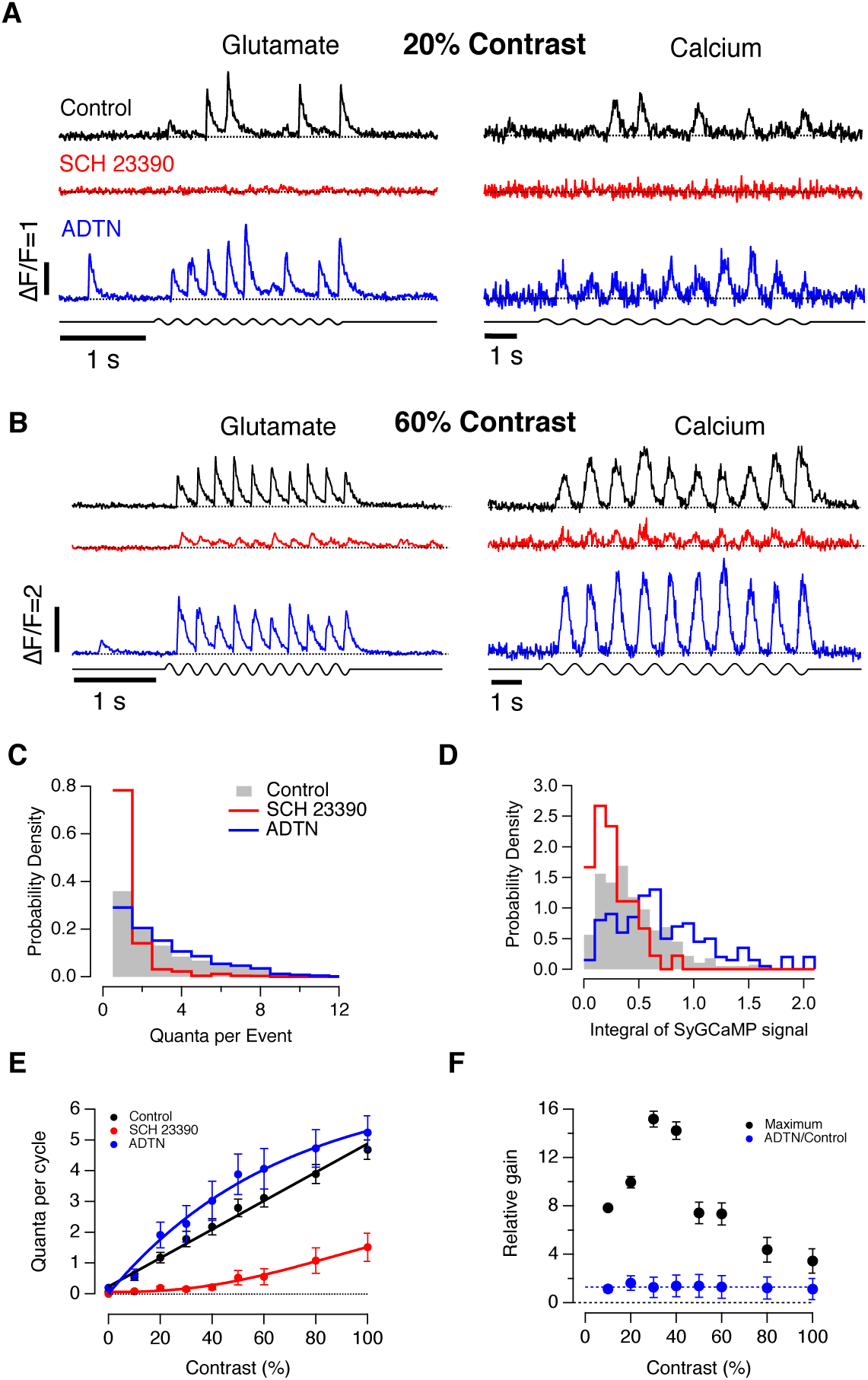
Endogenous dopamine potentiates presynaptic calcium transients and multivesicular release. **A.** Left panel: Examples of iGluSnFR signals from individual synapses elicited using a stimulus of 20% contrast (5 Hz) in three conditions: control (black), after intravitreal injection of SCH 23390 (red) and after injection of ADTN (blue). In these examples, SCH 23390 completely blocked synaptic events while ADTN increased both frequency and amplitude. Right panel: SyGCaMP6f signals from individual terminals elicited at 20% contrast (1 Hz). Note the lack of events in presence of SCH 23390. **B.** Left panel: Examples of iGluSnFR signals from individual synapses elicited using a stimulus of 60% contrast (5 Hz). Note the smaller glutamate transients with SCH 23390. Right panel: SyGCaMP6f signals from individual terminals at 60% contrast (1 Hz). **C.** Changes in the distribution of Q_e_ elicited at 60% contrast (5 Hz; control n = 37; ADTN n = 12; SCH 23390 n = 10). Statistical differences were present in control vs SCH 23390 (p<0.001, Chi-squared test). **D.** Changes in the distribution of the amplitude of calcium transients elicited at 60% contrast (1 Hz; control n = 450; ADTN n = 203; SCH 23390 n = 93). SCH 23390 shifted the distribution toward calcium transients of lower amplitude and ADTN to higher amplitudes (p<0.001, Wilcoxon rank test). **E.** Average contrast-response functions of bipolar cell synapses where the response (*R*) was quantified as the average of the number of quanta per cycle (Q_c_). Each point shows the mean ± s.e.m. (control, n=88 synapses; ADTN, n = 24; SCH 23390, n = 18). **F.** Relative changes in synaptic gain mediated by dopaminergic signalling as a function of stimulus contrast. The maximum change in gain (black) was calculated as the ratio of Q_c_ measured after injection of ADTN with respect to SCH 23390. The average quantal content of responses following injection of ADTN (blue) were 1.31 ± 0.06 that of control (dashed line) indicating that maximal activation of D1 receptors caused only a small increase in gain above the effect of endogenous dopamine levels.

Dopamine did not control the gain of synaptic transmission by changing the frequency of all synaptic symbols by the same factor. Instead, dopamine altered the distribution of symbols by increasing the proportion of larger multivesicular events encoding a given contrast. Examples of changes in the distribution of Q_e_ measured at 60% and 20% contrast are shown in Fig. 2C and Supplementary Figure 5, respectively. When D1 receptors were blocked 71 ± 6 % of stimulus cycles failed to elicit glutamate release at 60% contrast and only 8% of synaptic events were composed of 3 or more vesicles. But when D1 receptors were over-activated, only 23 ± 5% of stimulus cycles failed and 51% of events comprised 3 or more vesicles. Below we demonstrate how this shift towards larger synaptic events impacts on the temporal precision and efficiency of synaptic transmission (Figs. 5 and 6).

Changes in the distribution of event amplitudes (Fig. 2C) were paralleled by changes in calcium transients (Fig. 2D). For instance, activating D1 receptors also reduced the fraction of stimulus cycles failing to elicit a significant calcium influx and shifted the distribution of amplitudes toward higher values. While 11% of calcium transients had integral values of 0.5 or more with D1 receptors blocked, this increased to 67% when they were over-activated. These results are again consistent with the idea that the amplitude of an MVR event depends on the amplitude of the presynaptic calcium transient, with fluctuations in calcium influx as a major source of variability in synaptic output.

The effects of manipulating dopamine signalling across a range of stimulus contrasts are summarized in Fig. 2E and F. When synaptic output was quantified as Q_c_, the contrast-response function was almost perfectly linear under normal conditions (black trace in Fig. 2E) but when D1 receptors were blocked the relation became supralinear and contrasts up to ∼30% were barely signalled (red). The maximum change in gain caused by signalling through D1 receptors, calculated as the ratio of Q_c_ measured after injection of ADTN versus SCH 23390, is plotted as the black points in Fig. 2F. The range of modulation varied between 3.5-fold and 15-fold, depending on contrast, with the maximum at ∼30%. In contrast, antagonizing D2 receptors by injection of sulpiride (estimated final concentration of 0.4 μM) reduced synaptic gain by a factor of just 1.3 (Supplementary Figure 4). The observation that over-activating D1 receptors by injection of ADTN caused an increase in Q_c_ of only 30% relative to control (blue points in Fig. 2F), indicates that endogenous levels of dopamine were close to causing maximal potentiation of synaptic strength, at least under these experimental conditions.

### Substance P reduces the size of synaptic events by suppressing presynaptic calcium transients

Amacrine cells also release a number of neuropeptides, one of which is substance P (Yazulla and Studholme, 2001). Neurokinin-1 receptors activated by Substance P are expressed on terminals of some bipolar cells, and experiments in isolated neurons demonstrate that these act in a manner opposite to dopamine, *inhibiting* activation of L-type calcium channels (Ayoub and Matthews, 1992). Beyond this basic information, we have very little understanding about how Substance P might modulate retinal processing.

Examples of the effects of activating or blocking NK-1 receptors are shown in Fig. 3A and B. Substance P acted similarly on both ON and OFF synapses, so the two groups have been collected together (Supplementary Fig. 6). Injection of Substance P (estimated concentration of 200 nM) almost completely abolished transmission at contrasts up to 20% and this correlated with suppression of the presynaptic calcium transient (Fig. 3A-D). Conversely, the NK-1 receptor antagonist L-733060 (∼100 nM) increased both calcium influx and Q_c_. The degree to which Substance P acted on synaptic gain can be appreciated by comparing the average rate of vesicle release in response to a stimulus of 100% contrast: when NK1 receptors were antagonized release rate averaged 28 ± 2 vesicles/s falling to 9.8 ± 1.5 vesicles/s when Substance P was added.

**Figure 3.**
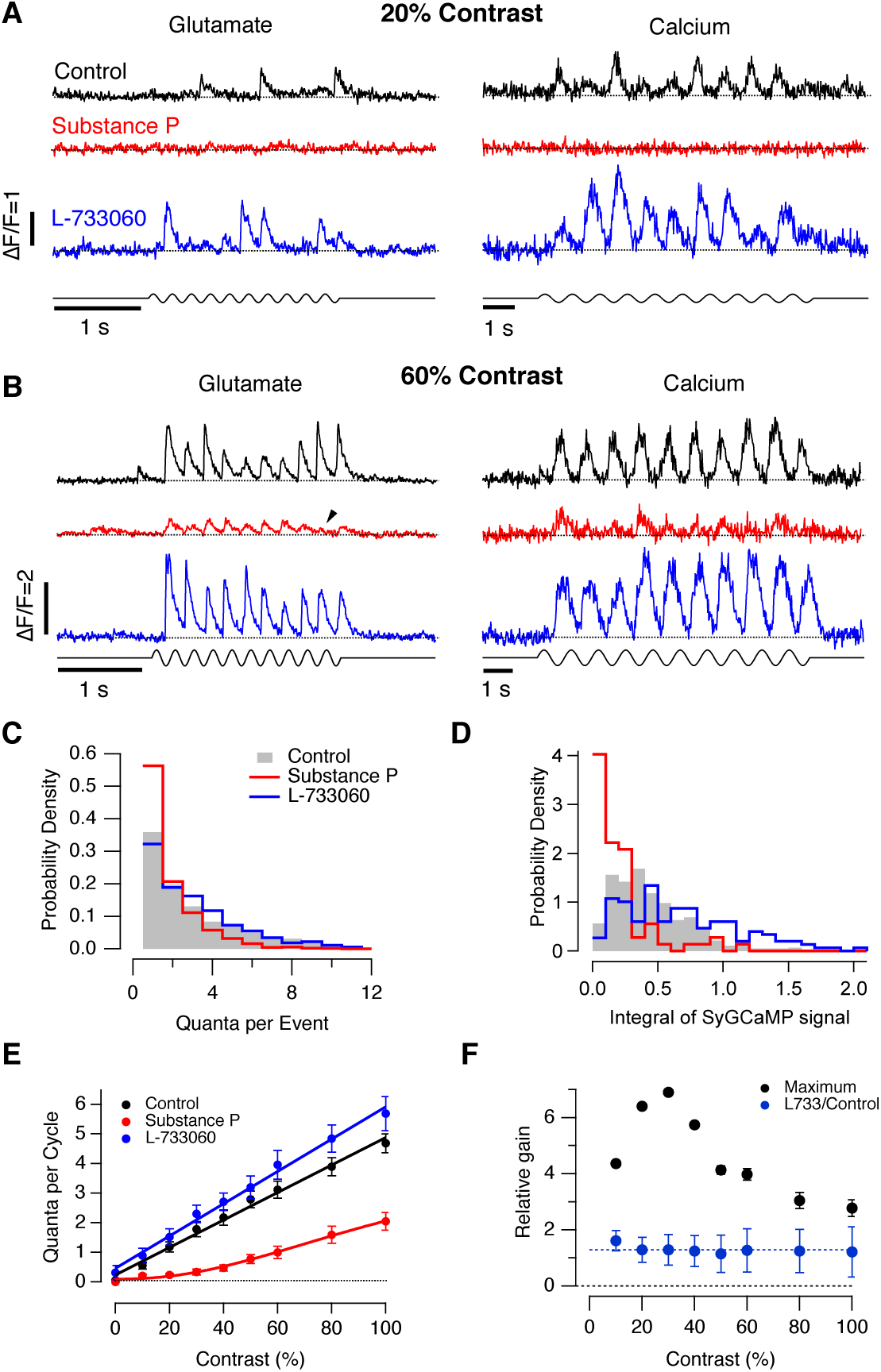
Substance P inhibits presynaptic calcium transients and multivesicular release. **A.** Left panel: Examples of iGluSnFR signals from individual synapses elicited using a stimulus of 20% contrast (5 Hz) in three conditions: control (black), after intravitreal injection of Substance P (red) and after injection of L-733060 (blue). In these examples, Substance P completely blocked synaptic events while L-733060 increased both frequency and amplitude. Right panel: SyGCaMP6f signals from individual terminals elicited at 20% contrast (1 Hz). Note the lack of events in presence of Substance P. **B.** Left panel: Examples of iGluSnFR signals from individual synapses elicited using a stimulus of 60% contrast (5 Hz). Note the smaller glutamate transients with injection of Substance P. Right panel: SyGCaMP6f signals from individual terminals at 60% contrast (1 Hz). **C.** Changes in the distribution of Q_e_ elicited at 60% contrast (5 Hz; control n = 37 synapses; Substance P, n = 14; L-733060, n=12). Substance P altered the distribution (p<0.001, Chi-squared test). **D.** Changes in the distribution of the amplitude of calcium transients elicited at 60% contrast (p<0.001, Wilcoxon rank test) (1 Hz; control, n = 450; Substance P, n = 73; L-733060, n = 149). Substance P shifted the distribution toward calcium transients of lower amplitude and L-733060 to higher amplitudes. **E.** Average contrast-response functions of bipolar cell synapses where the response (*R*) was quantified as the average of the number of quanta per cycle (Q_c_). Each point shows the mean ± s.e.m. (control, n = 88 synapses; Substance P, n = 24; L-733060, n = 22). **F.** Relative changes in synaptic gain mediated by neurokinin signalling as a function of stimulus contrast. The maximum change in gain (black) was calculated as the ratio of Q_c_ measured after injection of L-733060 with respect to Substance P. The ratio of L-733060 relative to control (blue) averaged 1.28 ± 0.04 (dashed line), indicating that endogenous Substance P was exerting close to the maximal possible action.

The effects of manipulating signalling through NK-1 receptors are summarized in Fig. 3E and F across a range of stimulus contrasts. Addition of substance P caused the contrast-response function to become supralinear and contrasts up to ∼20% were barely signalled (red). The maximum change in gain, calculated as the ratio of Q_c_ measured after injection of L-733060 versus Substance P, is plotted as the black points in Fig. 3F. The dynamic range varied between 3- and 7-fold, depending on contrast, with the maximum at ∼30%. Antagonizing the actions of endogenous Substance P by injection of L-733060 caused a 30% increase in Q_c_ relative to control conditions (blue points in Fig. 3F), indicating that the NK-1 signalling system was activated relatively mildly under these experimental conditions. These results demonstrate that Substance P decreases synaptic gain by suppressing the calcium transient driving transmission, thereby decreasing both the frequency and amplitude of synaptic events. More generally, results in Figs. 1-3, identify variability in presynaptic calcium signals as a major source of noise in transmission of the visual signal.

### Dopamine also acts downstream of the presynaptic calcium signal

Might dopamine or substance P also regulate transmission by acting on the presynaptic processes that are triggered by calcium influx? At the Purkinje cell to climbing fibre synapse, for instance, activation of PKA can potentiate MVR by increasing the number of vesicles docked and primed at the active zone (Vaden et al., 2019). To address this question, we assessed the effectiveness with which calcium released vesicles by measuring the relationship between Q_c_ and the Ca_c_ from records of the type shown in Fig. 1E and G (1 Hz stimulus). Not being able to make iGluSnFR and SyGCaMP6f measurements simultaneously, we instead manipulated the average amplitude of the presynaptic calcium transient by changing the contrast of the stimulus from 20% to 100% contrast. Note that measuring the relation between Q_c_ and Ca_c_ is not an attempt to assess the true calcium-dependence of the release process but rather an operational measure to detect changes in that relation.

Under normal conditions, the relation between Q_c_ and Ca_c_ was linear (open black circles in Fig. 4A), and it was not significantly affected when neurokinin signalling was increased by injection of Substance P (blue). But when D1 receptors were blocked by injection of SCH 23390, the relation became supralinear and at low stimulus contrasts calcium triggered release ∼10-fold less effectively, as can be seen more clearly in the log-log plot in Fig. 4B. Dopamine therefore increases the gain of transmission by at least two general mechanisms: potentiating the driver signal, calcium influx, and increasing the efficiency with which this signal triggers vesicle fusion, including multivesicular events.

**Figure 4.**
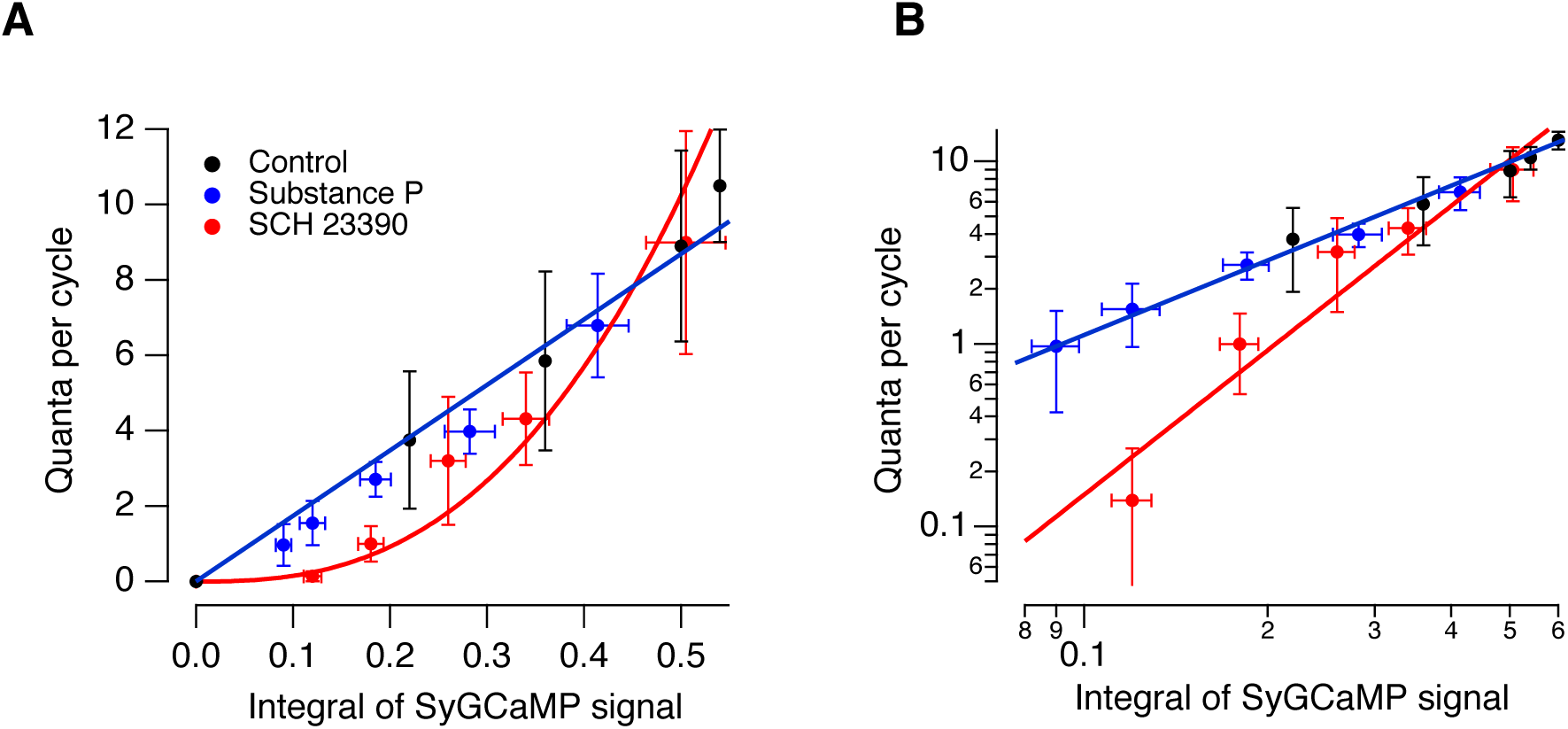
Dopamine modulates vesicle release downstream of the presynaptic calcium signal. **A.** Relationship between the number of vesicles released and the integral of SyGCaMP6f signal. Each point is the mean of each quantity at a different contrast (20%,40%,60%, 80% and 100%). The effects of NK-1 signalling were assessed by addition of Substance P (blue; n = 10 synapses), where there was no significant change in the relation relative to control (black; n = 8 synapses). In these conditions, the relation is described by a straight line through the origin with slope 14.4 ± 1.4. But after injection of the D1 receptor antagonist SCH 23390 (red; n = 9 synapses), the relation was best described as a power function with exponent 2.63 ± 0.4. **B.** Log-log plot of results in A. The different slopes of the lines through the points illustrate more clearly the higher power relation when D1 receptors were inactivated.

### Neuromodulators alter the efficiency of information transmission to RGCs

Multivesicular events of different amplitude can be considered as distinct symbols wthin the synaptic code and the information conveyed by each symbol can be evaluated using information theory (Meister and DeWeese, 1999). Using this approach, larger synaptic events were found to convey more information about the contrast of a visual stimulus compared to smaller events or individual vesicles, essentially because larger events are rare while being strongly correlated with higher contrasts (James et al., 2019). Further, multivesicular events have been shown to be more efficient in the sense of conveying more information *per vesicle*. How do neuromodulators that alter the distribution of these synaptic symbols affect information transmission?

To investigate this question we quantified the mutual information between a set of stimuli of varying contrasts and release events containing different numbers of quanta (Stone, 2018). The stimulus set consisted of 11 different contrasts over a range of ±10% around C_1/2,_ the contrast generating a half-maximal response, where contrast sensitivity is highest. Stimuli lasting 2 s were applied in a pseudorandom order and Figs. 5A and B compare signals from two active zones before and after injection of Substance P and ADTN, respectively. The time series of quantal events was divided into time bins of 20 ms, such that each bin contained either zero events or one event of an integer amplitude, from which we obtained the conditional distribution of Q given S, *p(Q|S)*, and the mutual information, *I(S;Q)*, the average amount of information conveyed across all possible event types. As expected, manoeuvres that reduced the average rate of vesicle release also reduced information transmission, Substance P by ∼42% (Fig. 5C; p<0.007, t-test) and SCH 23390 by ∼40% (Fig. 5D; p<0.002, t-test). Increasing activation of D1 receptors by injection of ADTN caused a small relative increase in the rate of vesicle release (Fig. 2E and F) but had no significant effect on mutual information (Fig. 5E).

**Figure 5.**
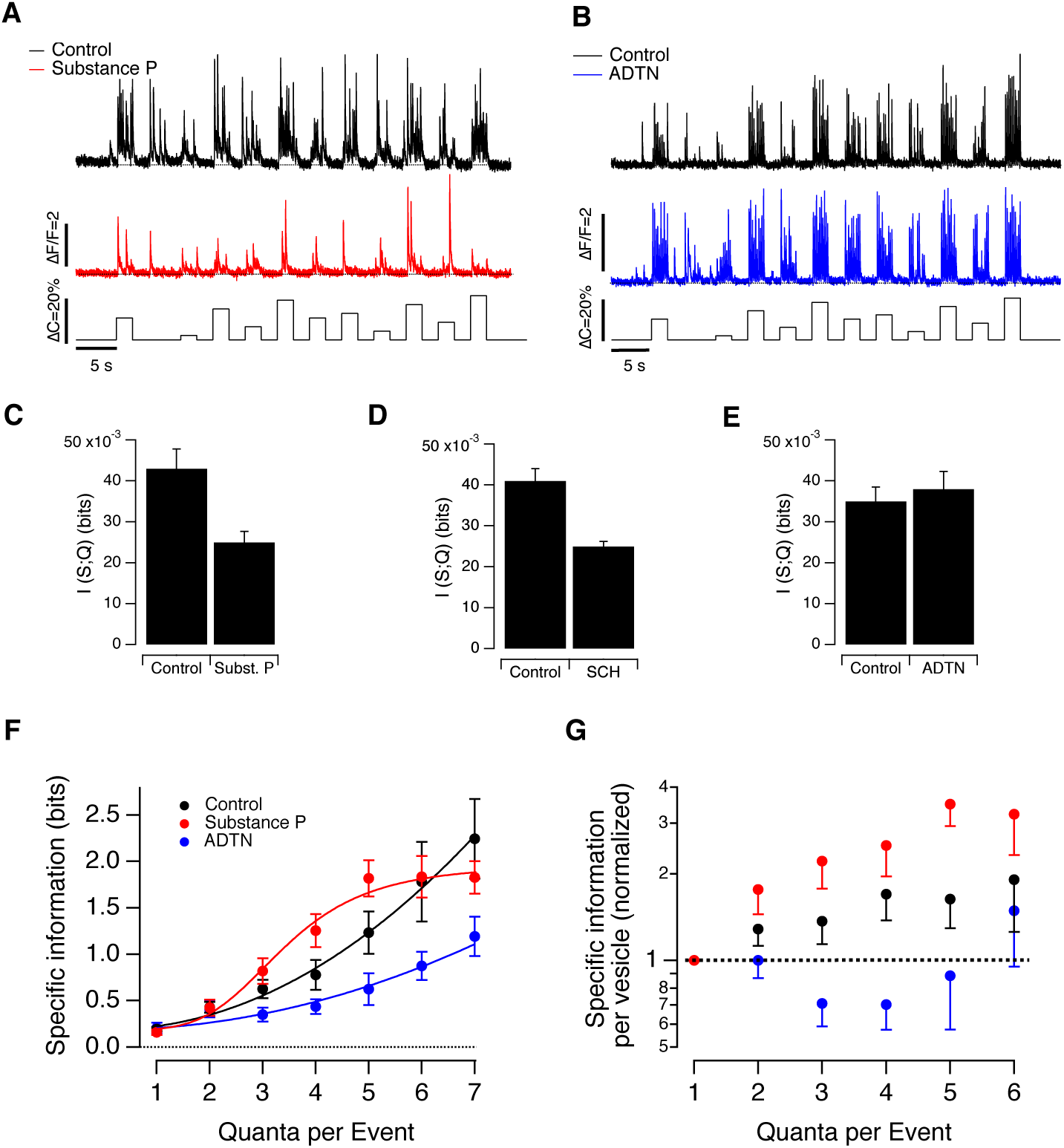
Neuromodulators alter the efficiency of information transmission to RGCs. **A.** Examples of synaptic responses over 12 different contrasts spanning ±10% around the contrast eliciting the half-maximal response (*C*_1/2_). Each step lasted 2 s (5 Hz modulation) and they were presented in two different pseudo-random orders, of which one shown. The distribution of this set of contrasts was uniform. Note the decrease in frequency and amplitude of glutamate transients in Substance P relative to control. **B.** Further examples showing the increased amplitude and frequency of events after injecting ADTN. **C.** Estimation of mutual information I (S;Q) before and after intravitreal injection of Substance P (paired measurements in the same synapses; n = 13). Substance P reduced the mutual information values 42% (p<0.007; t-test). **D.** Estimation of I(S;Q) after administration of SCH 23390. Mutual information was reduced by 40% (p<0.002, t-test). **E**. Estimation of I(S;Q) after administration of ADTN. Mutual information was not significantly affected (p<0.48, t-test). **F.** Specific information per event (*I*_*2*_, bits) as a function of *Qe*, the number of vesicles comprising the event. The curve describing the control (black; n = 27 synapses) is a power function of the form *i* = *i*_*min*_ + *AQ*_*e*_^*x*^, with *i*_*min*_ = 0.196 bits, *A* = 0.04 ± 0.02 and *x* = 2.05 ± 0.22. After injection of ADTN (blue, n = 13 synapses), A = 0.02 ± 0.014 and x =1.99 ± 0.42, with *i*_*min*_ unchanged. Events composed of 4-7 quanta transferred less information compared to control (p<0.05; t-test). When synaptic gain was reduced by injection of Substance P (red; n = 14 synapses), this relation was better described by a Hill function of the form *i* = *i*_*min*_ + (*A-i*_*min*_*)Q*_*e*_^*x*^ /(*Q*_*e*_^*x*^ *+ C*^*x*^) where A =1.97 ± 0.22, C =3.42 ± 0.36 and x = 4.14 ± 1.0 (again with *i*_*min*_ unchanged). Events composed of 4-5 quanta transferred more information relative to control (p<0.05; t-test). **G.** Specific information per vesicle normalized to the value measured for a uniquantal event in the same synapse (*i*′). Events composed of 2-5 quanta were transferred information more efficiently when synaptic gain was reduced by Substance P compared to when it was increased by activating D1 receptors with ADTN (p<0.004; t-test).

Changes in total information transmission did not, however, scale in a simple way with changes in the average rate of vesicle release. The relation between specific information (*I*_*2*_, bits) and *Q*_e_, the number of vesicles comprising the event, is shown in Fig. 5F. Under control conditions and after injection of ADTN, the relation could be described as a power function with an exponent of ∼2 (black and blue traces). Notably, the amount of information carried by vesicles released individually was relatively constant across all conditions, averaging 0.19 bits, but events composed of 4-6 vesicles transmitted significantly less information when synaptic gain was increased by increased activation of D1 receptors. Neurokinin signalling, which reduced synaptic gain, exerted the opposite action, *increasing* the information content of events composed of 4-5 quanta (Fig. 5F).

To evaluate these measurements in terms of ‘vesicle efficiency’, we divided the amount of information transmitted by each event type by the number of vesicles it contained. The change in this quantity as a function of Q_e_ is plotted in Fig. 5G, normalized to the value measured in the same synapse for vesicles released individually. Substance P (red points) increased the specific information per vesicle relative to control (black) by factors varying between 40% and 110%, depending on Q_e_. Activation of D1 receptors by ADTN had the opposite effect, most significantly for events comprising 3-4 vesicles (blue). These results reveal that neuromodulatory increases in synaptic gain come at the cost of less efficient use of vesicles for transmitting information.

### Neuromodulators regulate the temporal precision of the vesicle code

Multivesicular release also impacts on the temporal precision with which ribbon synapses transmit sensory information. In ribbon synapses of retinal bipolar cells and auditory hair cells, the larger the exocytic event, the more precisely it is synchronized to a stimulus (Li et al., 2014; James et al., 2019). This property of MVR is signficant in the context of vision because larger events are more common at higher contrasts (Figs. 1-3) when the spike output of post-synaptic ganglion cells also becomes more precise in time (Berry et al., 1997). How do neuromodulators that regulate MVR affect the temporal precision of synaptic transmission?

The timing of synaptic events were measured relative to the phase of a 5 Hz stimulus of 60% contrast. The traces in Fig. 6A show examples of iGluSnFR responses measured under control conditions and two extremes: potentiation of synaptic strength by injection of ADTN and suppression of transmission by injection of Substance P. The arrowheads highlight events occurring at different phases and numbers represent the number of quanta released within the cycle of stimulation. Under normal conditions, individual vesicles were released with a standard deviation (‘temporal jitter’) of 27 ± 2 ms (15 synapses) which was not significantly altered by neuromodulators (Fig. 6B). The larger the multivesicular event the less it varied in time, but while Substance P caused a precision of ∼11 ms to be achieved with events comprising just two quanta (red points), activation of D1 receptors caused this precision to be achieved only with events comprising 4-5 vesicles. A decrease in synaptic strength and MVR was therefore associated with an increase in the temporal precision with which the visual signal was transmitted to the inner retina.

**Figure 6.**
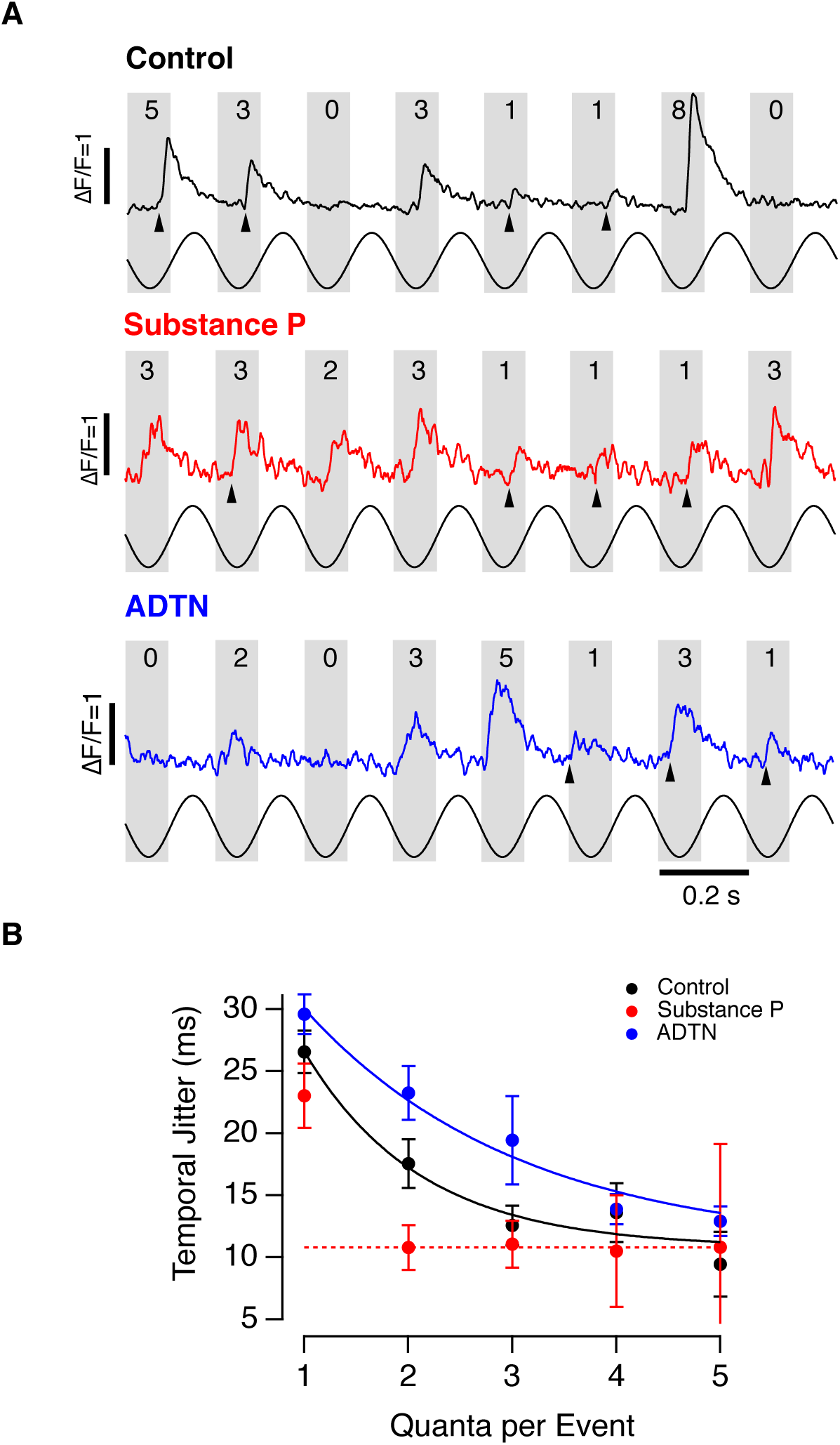
Neuromodulators regulate the temporal precision of multivesicular events. Example recordings from synapses stimulated at 60% contrast under control conditions (black, top), after injection of Substance P (red, middle) or ADTN (blue, bottom). The modulation in intensity (5 Hz sinewave) is shown below. Arrowheads highlight events occurring at different phases of the stimulus and numbers indicate estimated number of quanta in each event. **B.** Temporal jitter of events composed of different numbers of quanta (*Q*_*e*_) measured from the experiments in A (control, n = 12 synapses; Substance P, n = 10; ADTN, n = 12). Neuromodulators had no significant effect on release of individual vesicles. In the presence of Substance P, events composed of 2 or more quanta jittered by an average of 10.8 ms (red dashed line). The solid lines describing this relation under control conditions and in ADTN are exponentials that decay to this same limiting value as eQe/x, where x = 0.74 ± 0.33 under control conditions and 1.60 ± 0.31 in ADTN (mean ± sd). This difference is significant at p <0.003 (t-test). Thus, the jitter in timing of multivesicular events was lower when synaptic gain was reduced by Substance P and higher when it was increased by activating D1 receptors.

### Neuromodulators alter the synaptic transfer function

Manipulations of dopamine or Substance P signalling that caused a decrease in the frequency of synaptic events also reduced the average rate of information transmission (Fig. 5C and D). Framed in terms of signal-detection theory, a reduction in the average rate of synaptic events will also increase the duration of the time-window required to detect a change in activity, making the retina less sensitive to brief stimuli (James et al., 2019). We therefore wondered whether neuromodulatory effects that reduced synaptic gain also tuned the output of bipolar cells to lower temporal frequencies and found that they did.

To measure the synaptic transfer function we applied a full-field stimulus modulated at frequencies between 0.5 and 30 Hz (60% contrast). Example responses are shown in Fig. 7A for control (black trace), Substance P (red) and ADTN (blue). Under control conditions, the cut-off frequency (f_c_) at -3 dB of the maximum response was 9.6 Hz (42 synapses). Enhancing activation of D1 receptors by injection of ADTN increased synaptic gain across all frequencies, with a maximum of 3.2-fold at 10 Hz, while also shifting f_c_ slightly to 11.1 Hz (Fig. 7C, 12 synapses). Addition of Substance P decreased synaptic gain by a maximum factor of 6.2-fold at a frequency of 8 Hz, while shifting f_c_ to 4 Hz (n = 10 synapses). Notably, Substance P also altered the transfer characteristics from band-pass to low-pass (Fig.7B). Although these experiments do not determine the loci at which Substance P exerts these effects, they demonstrate that changes in synaptic gain occur in parallel with changes in the frequency response of the retinal circuit.

**Figure 7.**
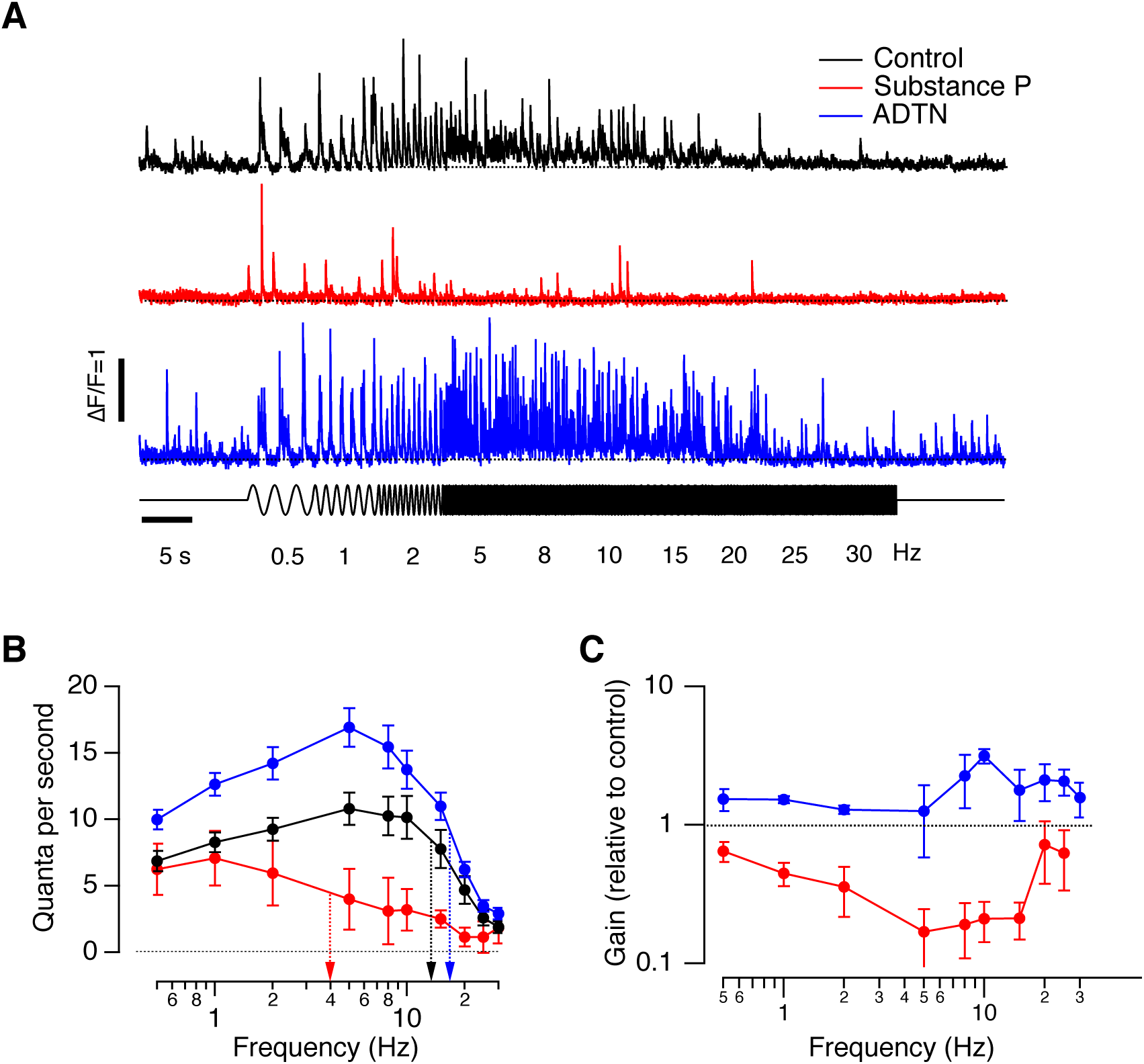
Neuromodulators regulate temporal filtering of visual stimuli. **A.** Example of iGluSnFR signals at individual synapses stimulated at a range of frequencies from 0.5 to 30 Hz in control conditions (top, black), and after intravitreal injection of Substance P (red, middle) or ADTN (blue). Lower trace is the stimulus (60% contrast, sinewave). **B.** Synaptic transfer function measured as release rate (quanta per second) as a function of frequency from experiments of type shown in A. Dotted lines represent the average cut-off frequencies (f_c_) at -3 dB of the maximum response. F_c_ values were 9.6 Hz, 4.0 Hz and 11.1 Hz for control, Substance P and ADTN conditions, respectively. Note that the transfer function is bandpass under control conditions and when synaptic gain is increased by ADTN, but shifts to low-pass when gain is reduced by Substance P. Control, n=42 synapses; Substance P, n =10; ADTN, n=13). **C.** Release rate relative to control as a function of frequency (log-log plot). Dashed line indicates no change. The maximal reduction in gain caused by Substance P occurred over the frequency range 5-15 Hz.

## Discussion

The ability to count vesicles released at individual active zones of a live animal has allowed us to investigate the mechanisms by which neuromodulators act on information transmission across a synapse, much as the ability to count spikes has allowed investigation of the spike code operating within neurons (Berry et al., 1997; Gollisch and Meister, 2008). Most fundamentally, this study demonstrates that individual synapses transmit the visual signal with a high degree of variability, in large part reflecting fluctuations in the processes controlling calcium influx into the presynaptic compartment (Figs. 1-3). Dopamine and Substance P act on calcium influx to alter both the average rate and amplitude of synaptic events (Figs. 2-3), while dopamine additionally enhances gain at low contrasts by increasing the number of vesicles released by smaller calcium signals (Fig. 4). By acting on the process of multivesicular release, Substance P and dopamine regulate the efficiency with which vesicles code information (Fig. 5) as well as the temporal precision of transmission (Fig. 6). These modulations in gain were strongly frequency-dependent, and neurokinin signalling shifted the transfer characteristics of the retinal circuit from band-pass to low-pass (Fig. 7).

### Mechanisms regulating multivesicular release

What are the electrical events that convert the current flowing down the axon of the bipolar cells into multivesicular release? Capacitance measurements in isolated bipolar cells have previously shown that, when strongly depolarized, each active zone can release several vesicles within milliseconds (Mennerick and Matthews, 1996; Neves and Lagnado, 1999; Burrone and Lagnado, 2000). It was less clear whether this occurred during normal visual processing because voltage measurements in the soma of bipolar cells demonstrate that this compartment is depolarized passively and by no more than ∼10 mV (Slaughter and Miller, 1981). We now understand, through electrophysiology and imaging, that the terminal of bipolar cells is in fact an active compartment, in which voltage-dependent calcium channels and calcium-activated potassium channels generate voltage spikes (Burrone and Lagnado, 1997). Consistent with this idea, dopamine and Substance P, which both alter the voltage threshold for activation of L-type calcium channels (Ayoub and Matthews, 1992; Heidelberger and Matthews, 1994) also altered the frequency and amplitude of the presynaptic calcium transient triggered by a periodic stimulus, thereby altering the distribution of MVR events (Figs. 1-3).

What factors might make the presynaptic depolarization so variable from trial-to-trial? Spike generation in bipolar cells can be turned and off by changes in membrane potential as small as 2-3 mV and will therefore be sensitive to electrical noise (Baden et al., 2011). The stochastic opening and closing of voltage-dependent ion channels has long been known to affect the consistency of neuronal responses (White et al., 2000) and it is notable that the calcium-activated potassium channels in the terminal are of large conductance (Burrone and Lagnado, 1997), a type which have been shown to determine the variability of calcium spikes in neuroendocrine cells (Richards et al., 2020). An additional, extrinsic, source of voltage fluctuations is likely to be GABAergic amacrine cells that synapse directly onto the terminal and show high rates of spontaneous release (Sagdullaev et al., 2011; Palmer, 2006). A more quantitative understanding of these sources of electrical noise will be important in building a more complete picture of the mechanisms that control the reliability of ribbon synapses as they transmit the visual signal.

A related question is how much additional noise is contributed by the stochasticity of the molecular processes subsequently triggered by calcium influx. Electrical recordings from the mouse retina indicate that rod-driven bipolar cells display almost perfect correlations in glutamate release across different output synapses in the dark, which then breaks down as light levels increase (Grimes et al., 2014). In contrast, the retina of zebrafish is cone-dominated for operation under daylight conditions and monitoring iGluSnFR signals from different active zones in the same terminal demonstrate that the stochasticity of the release process makes a major contribution to variability in synaptic output (James et al., 2019). Separately quantifying the contributions of electrical and molecular noise will require simultaneous recording of glutamate release from different active zones in tandem with either voltage or calcium recordings in the presynaptic compartment. Electrophysiology does not allow one to isolate the output of a single active zone, but the optical approach we describe here might make this experiment possible if iGluSnFR could be imaged together with voltage or calcium reporters emitting at different wavelengths and with sufficiently high signal-to-noise ratio.

### Dopamine and Substance P as neuromodulators in retina

Dopamine release from amacrine cells varies according to a number of factors, beginning with a circadian cycle in which high levels occur during the day and low levels at night (Doyle et al., 2002). A second level of control is exerted by light directly, with dopamine released under brighter conditions adapting the retina to high-acuity vision. A third mechanism of regulation derives from the sense of smell, with dopamine release being stimulated through the olfacto-retinal pathway that is activated by different chemical stimuli, including food. Odorants effectively trigger changes in brain state, adjusting visual processing in the retina in tandem with behaviours ranging from visual escape responses to mating (Dowling, 2013; Esposti et al., 2013; Li and Dowling, 2000; Maaswinkel and Li, 2003). It will be important to understand how these various processes interact in regulating retinal computations in relation to different visually-driven behaviours. Although they all engage dopmaine signalling, we do not know if they alter transmission of the visual signal in the same manner.

Dopamine acts on a number of voltage-gated ion channels, especially those sensitive to cAMP metabolism, such as the HCN channel found in terminals of photoreceptors and bipolar cells (Muller et al., 2003). Signal flow is also adjusted by closing gap junctions that couple horizontal cells, rods and cones and some bipolar and amacrine cells (Jackson et al., 2012; Roy and Field, 2019). The specific actions of dopamine depend on which of the two major subtype of dopamine receptor a neuron expresses. Modulation of transmission from bipolar cells was mediated through the D1 receptor class located on synaptic terminals of bipolar cells (Fig. 2), while D2 receptors had relatively little effect (Supplemtary Figure 4). But the D1 receptors are also found on horizontal cells and some amacrine cells (Kothmann et al., 2009; Xin and Bloomfield, 1999) and it may be that changes in contrast-response function reflect both direct actions on the terminal and indirect actions through other components of the retinal circuit.

The decrease in synaptic gain caused by Substance P was, at least qualitatively, antagonistic to the effects of dopamine, even though both were active simultaneously. Our analysis revealed, however, important differences in the mechanism by which these modulators acted on the synapse and the consequences for coding the visual signal. While Substance P reduced synaptic gain simply through modulation of calcium influx, dopamine additionally increased the efficiency with which calcium triggered release (Fig. 4). Substance P also increased the efficiency and temporal precision of synaptic transmission by altering the frequency and amplitude distribution of synaptic events (Fig. 5 and 6), as well as retuning the retinal circuit to lower temporal frequencies (Fig. 7). Placing these various effects in a broader context will require a better understanding of the factors that regulate the release of Substance P and other neuromodulators and their consequences on visually-driven behaviours. This study has demonstrated that changes in the vesicle code are a fundamental mechanism by which such neuromodulators adapt the retinal circuit for operation under different conditions.

## Methods

### Zebrafish husbandry

Fish were raised and maintained under standard conditions on a 14 h light/10 h dark cycle (Odermatt et al., 2012). To aid imaging, fish were heterozygous or homozygous for the casper mutation which results in hypopigmentation and they were additionally treated with1-phenyl-2-thiourea (200 µM final concentration; Sigma) from 10 hours post fertilization (hpf) to reduce pigmentation. All animal procedures were performed in accordance with the Animal Act 1986 and the UK Home Office guidelines and with the approval of the University of Sussex Animal Welfare and Ethical Review Board. More information about experimental design and reagents is available in the Life Sciences reporting Summary.

### Transgenic fish

*Tg(–1.8ctbp2:Gal4VP16_BH)* fish that drive the expression of the transcriptional activator protein Gal4VP16 were generated by co-injection of I-SceI meganuclease and endofree purified plasmid into wild-type zebrafish with a mixed genetic background. A myocardium-specific promoter that drives the expression of mCherry protein was additionally cloned into the plasmid to allow for phenotypical screening of founder fish. *Tg(10xUAS:iGluSnFR_MH)* fish driving the expression of the glutamate sensor iGluSnFR under the regulatory control of the 10 x UAS enhancer elements were generated by co-injection of purified plasmid and tol2 transposase RNA into offspring of AB wildtype fish outcrossed to casper wildtype fish. The sequences for the myocardium-specific promoter driving the expression of enhanced green fluorescent protein (mossy heart) were added to the plasmid to facilitate the screening process. *Tg(–1.8ctbp2:SyGCaMP6)* fish were generated by co-injection of I-SceI meganuclease and endofree purified plasmid into wild-type zebrafish with a mixed genetic background (Thermes *et al*. <u>2002</u>). The GCaMP6f variant (alternative name GCaMP3 variant 10.500) was kindly provided by L. Looger (Janelia Farm). This variant holds a T383S mutation in comparison to the commercially available GCaMP6-fast version (Addgene plasmid 40755).

### Multiphoton Imaging *In Vivo*

Experiments were carried out in a total number of 132 zebrafish larvae (7–9 days post-fertilization). Fish were immobilized in 3% low melting point agarose (Biogene) in E2 medium on a glass coverslip (0 thickness) and mounted in a chamber where they were superfused with E2. Imaging was carried out using a two-photon microscope (Scientifica) equipped with a mode-locked titanium-sapphire laser (Chameleon, Coherent) tuned to 915 nm and an Olympus XLUMPlanFI 20x water immersion objective (NA 0.95). To prevent eye movements, the ocular muscles were paralyzed by injection of 1 nL of α-bungarotoxin (2 mg/mL) behind the eye. The signal-to-noise ratio for imaging was optimized by collecting photons through both the objective and a sub-stage oil condenser (Olympus, NA 1.4). Emission was filtered through GFP filters (HQ 535/50, Chroma Technology) before detection with GaAsP photomultipliers (H7422P-40, Hamamatsu). The signal from each detector passed through a current-to-voltage converter and then the two signals were added by a summing amplifier before digitization. Scanning and image acquisition were controlled under ScanImage v.3.6 software (Pologruto et al., 2003). In iGluSnFR recordings images were acquired at 10 Hz (128 × 100 pixels per frame, 1 ms per line) while linescans were acquired at 1 kHz. In calcium recordings images were acquired at 50 Hz (128 × 20 pixels per frame, 1 ms per line).

Full-field light stimuli were generated by an amber LED (lmax = 590 nm, Thorlabs), filtered through a 590/10 nm BP filter (Thorlabs), and delivered through a light guide placed close to the eye of the fish. Stimuli were normally delivered as modulations around a mean intensity of ∼165 nW/mm^2^ and the microscope was synchronized to visual stimulation.

### Drug injections

To manipulate Substance P and Dopamine signalling *in vivo*, we injected agonist and antagonists of their receptors into the anterior chamber of the retina. Substance P was manipulated by injecting the agonist of the NK-1 receptor Substance P (Tocris) to an estimated final concentration of 200 nM, and the effects of endogenous Substance P were antagonized by injection of L-733060 (Tocris) at estimated final concentration of 100 nM. Dopamine signalling was manipulated by injecting the antagonist of D1 receptors SCH 23390 at a final estimated concentration of 1 μM (Sigma), the selective D2 receptor antagonist Sulpiride (Sigma) was injected at a final estimated concentration of 400 nM. Finally, the long-lasting dopamine receptor ligand [3H] 2-amino-6,7-dihydroxy 1,2,3,4-tetrahydronapthalene (ADTN) (Sigma) was injected to a final estimated concentration of 200 nM. We confirmed that these drugs gained access by including 1 mM Alexa 594 in the injection needle; within 5 mins of injection the dye could be detected within the inner plexiform layer of the retina. Vehicle injection did not affect synaptic responses to varying contrast. The effects of dopamine and Substance P are dependent on the circadian cycle, so all experiments were carried out at 2-6 pm (Doyle et al., 2002).

### Statistics

All data are given as mean ± s.e.m. unless otherwise stated in the figure legends. All statistical tests met appropriate assumptions and were calculated using inbuilt functions in IgorPro (Wavemetrics). When data were not normally distributed we used non-parametric tests. All tests were two-sided and significance defined as *P* < 0.05. Data collection was not randomized because all experiments were carried out within one set of animals. Delivery of different stimuli was randomized where appropriate. Data were only excluded from the analysis if the signal-to-noise ratio (SNR) of the iGluSnFR signals elicited at a given synapse was not sufficient to detect unitary responses to visual stimuli with a SNR of at least three.

### Calculation of temporal jitter

In order to quantify variability in the timing of glutamatergic events, we first calculated the vector strength, r_q_, for events composed of q quanta:

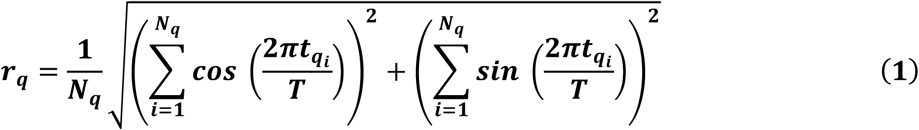

where t_qi_ is the time of the i^th^ q-quantal event, T is the stimulus period, and N_q_ is the total number of events of composed of q-quanta. The temporal jitter, J_q_, can then be calculated as:

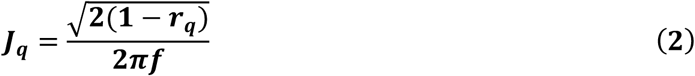

where f is the stimulus frequency.

### Calculations based on Information Theory

To quantify the amount of information about a visual stimulus that is contained within the sequence of release events from an active zone we first needed to convert bipolar cell outputs into a probabilistic framework from which we could evaluate the specific information (I_2_), a metric that quantifies how much information about one random variable is conveyed by the observation a specific symbol of another random variable (Stone, 2018). The time series of quantal events was converted into a probability distribution by dividing into time bins of 20 ms, such that each bin contained either zero events or one event of an integer amplitude. We then counted the number of bins containing events of amplitude 1, or 2, or 3 etc. By dividing the number of bins of each type by the total number of bins for each different stimulus, we obtained the conditional distribution of **Q** given **S**, *p*(***Q***|***S***), where **Q** is the random variable representing the *quanta/bin* and **S** is the random variable representing the *stimulus contrasts* presented throughout the course of the experiment. In the absence of information about the distribution of contrasts normally experienced by a larval zebrafish, a uniform distribution was used. We then computed the joint probability distribution by the chain rule for probability (given the experimentally defined uniform distribution of stimuli **S**):

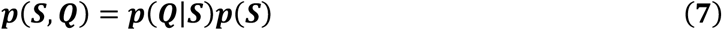

In order to convert this distribution into the conditional distribution of S given Q, we used the definition of the conditional distribution:

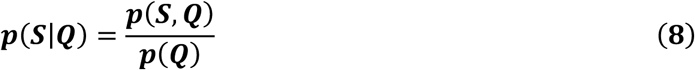

From these distributions we computed two metrics: the mutual information I(**S;Q**) (de Ruyter van Steveninck et al., 1997) and specific information I_2_(**S;q**) (Meister and DeWeese, 1999). Mutual information is defined traditionally as:

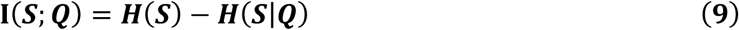

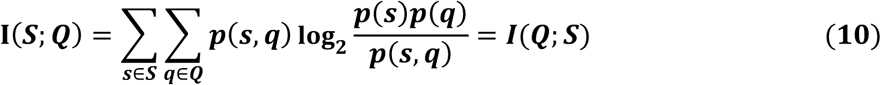

The specific information, I_2_(**S;q**), is defined as the difference between the entropy of the stimulus S minus the conditional entropy of the stimulus given the observed symbol in the response q:

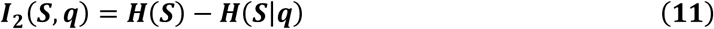

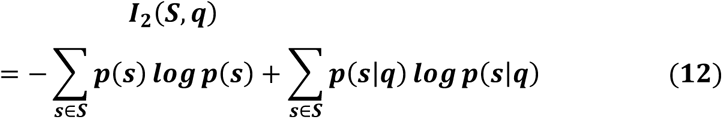

representing the amount of information observing each quantal event type q ϵ **Q** carries about the stimulus distribution **S**. Note that mutual information can also be computed from the specific information as the dot product of the specific information vector ***I***_**2**_ and the vector describing the probability of an event of a given quantal size ***p***(***q***). This adds to the interpretability of both metrics – the specific information is the amount of information a single (specific) symbol gives about the stimulus, and the mutual information is the average amount of information gained from observing any symbol about the stimulus.

Measuring entropy and mutual information from neural responses can be a challenging problem. Estimates require sampling from an unknown discrete probability distribution, and in many cases recording sufficient samples to observe all non-zero probability events is neither tractable nor practical. The biases introduced by undersampling can be a particular problem when the full support of the distribution (all values that map to non-zero probabilities) is high. Within the past few decades, various approaches to correcting biases in information theoretic analyses have been developed (Pola, 2003). However, as the distributions of interest in this work have both a small support and are well sampled, we have opted to use standard estimates for the quantities of interest.

## Acknowledgements

The authors express many thanks to all the members of Lagnado laboratory for discussion. We also thanks H. Smulders and N. Bashford for looking after the zebrafish. This work was supported by grants to L.L. from the Wellcome Trust (102905/Z/13/Z) and an EU International Training Network (H2020-MSCA-ITN-2015-674901).

## Author contribution

JM-D: Conceived and designed experiments, carried out 2-photon imaging experiments, analyzed data and helped to prepare the manuscript. BJ: carried out analysis and wrote code. LL: Conceived the project, designed experiments, analyzed data, wrote code and wrote the manuscript.

## Supplementary Figures

**Figure S1.**
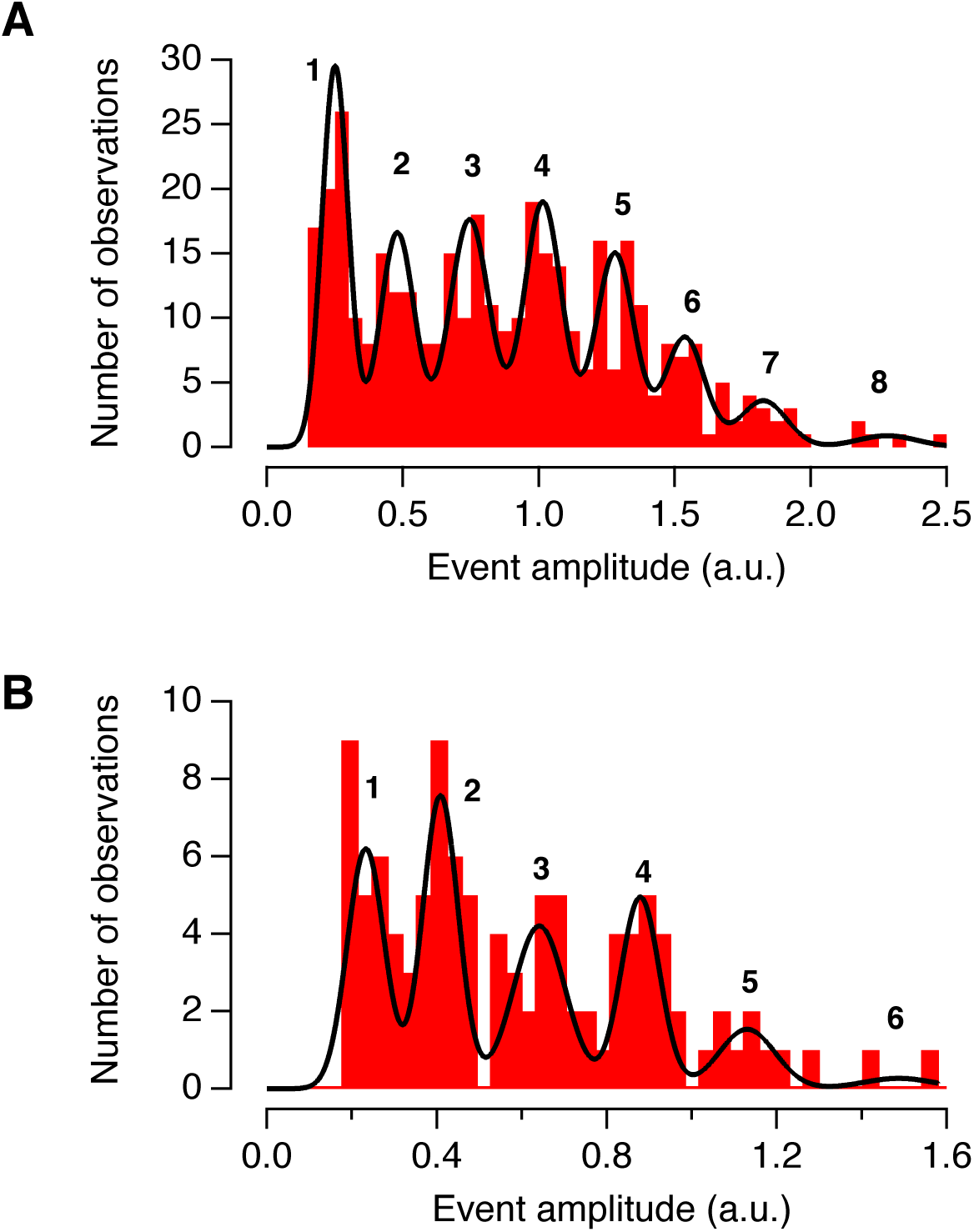
Estimating the iGluSnFR signal generated by a single vesicle. **A.** Histogram of event amplitudes for a representative active zone (373 events accumulated using stimulus contrasts of 20%, 60% and 100%). The black line is a fit of eight Gaussians, identified using a Gaussian mixture model. Note that the variance of successive Gaussians did not increase in proportion to the peak number. The first peak had a value of 0.24, and the distance between peaks averaged 0.25, indicating the existence of a quantal event equivalent to ∼0.25. **B.** A second example of an amplitude histogram measured at a single contrast (50%) from another active zone (106 events).

**Figure S2.**
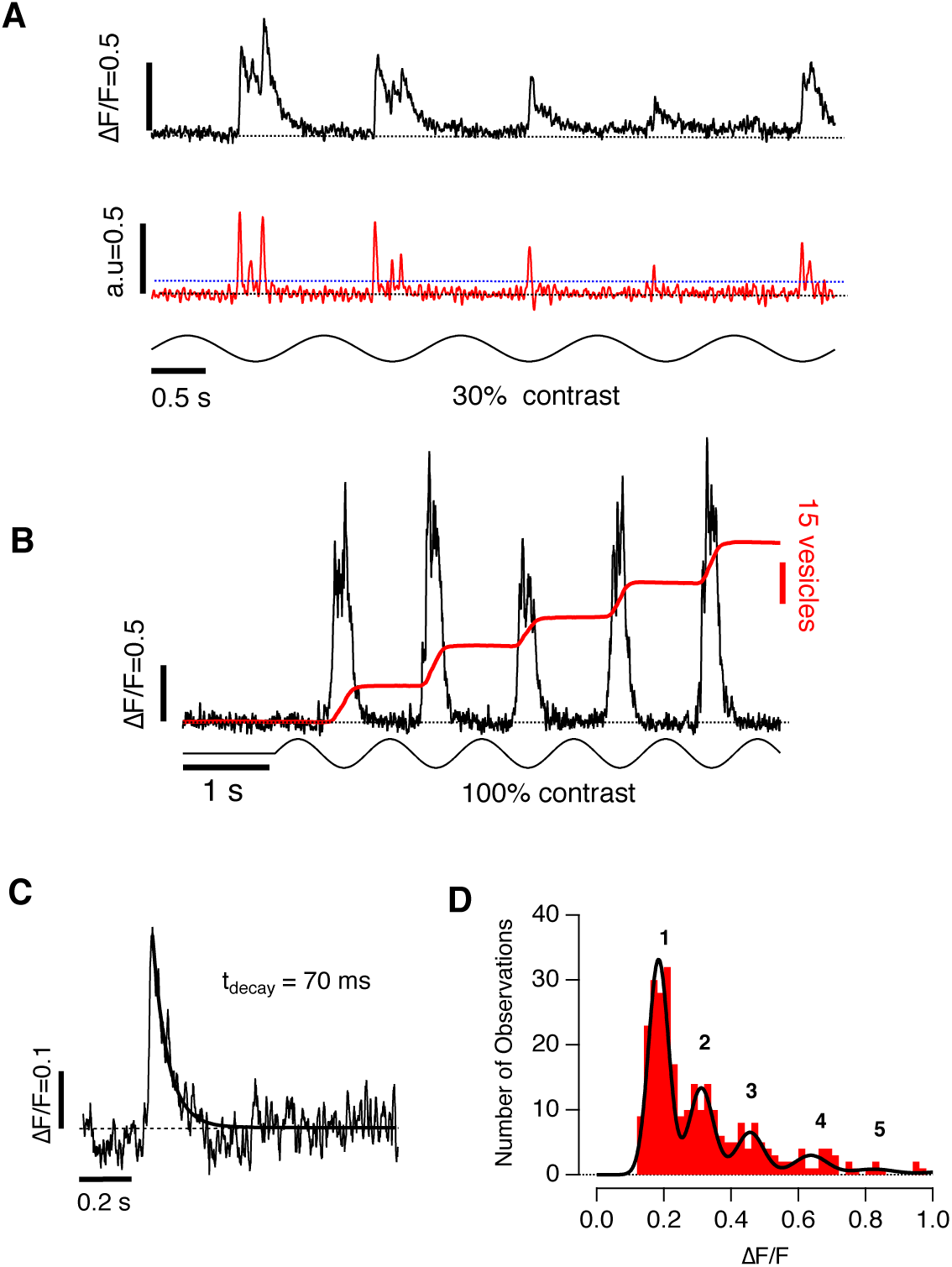
Quantifying vesicles released during large bursts of activity. **A.** Recordings from an individual active zone stimulated at 30% contrast and a frequency of 1 Hz. Individual release events were easily identified in the raw trace (top) and by deconvolution (bottom). The blue dashed line shows the threshold set for counting events. **B.** Example of a recording from an active zone stimulated at 100% contrast and 1 Hz, when individual synaptic events are not easy to distinguish. In these cases we did not attempt to obtain accurate timing information but instead measured the number of vesicles released in a burst by integration (red trace). The scaling factor for conversion of the integrated iGluSnFR signal into numbers of vesicles was the integral of the unitary event, calculated for each active zone as the product of the unitary event amplitude and decay time-constant. In this example, the time-constant of decay was 70 ms (C) and the unitary amplitude was 0.18 (D). **C.** Single exponential fit to a single iGluSnFR event from the synapse in B. **D.** Histogram of event amplitudes for 5 different active zones, including that featured in B (n = 290 events accumulated using stimulus contrasts of 20% and 30%; 1 Hz stimulus). The black line is a fit of Gaussians, identified using a Gaussian mixture model as in Fig. S1. In this set of experiments the first peak had a value of 0.18 and the distance between peaks averaged ∼0.18, indicating the existence of a quantal event equivalent to ∼0.18.

**Figure S3.**
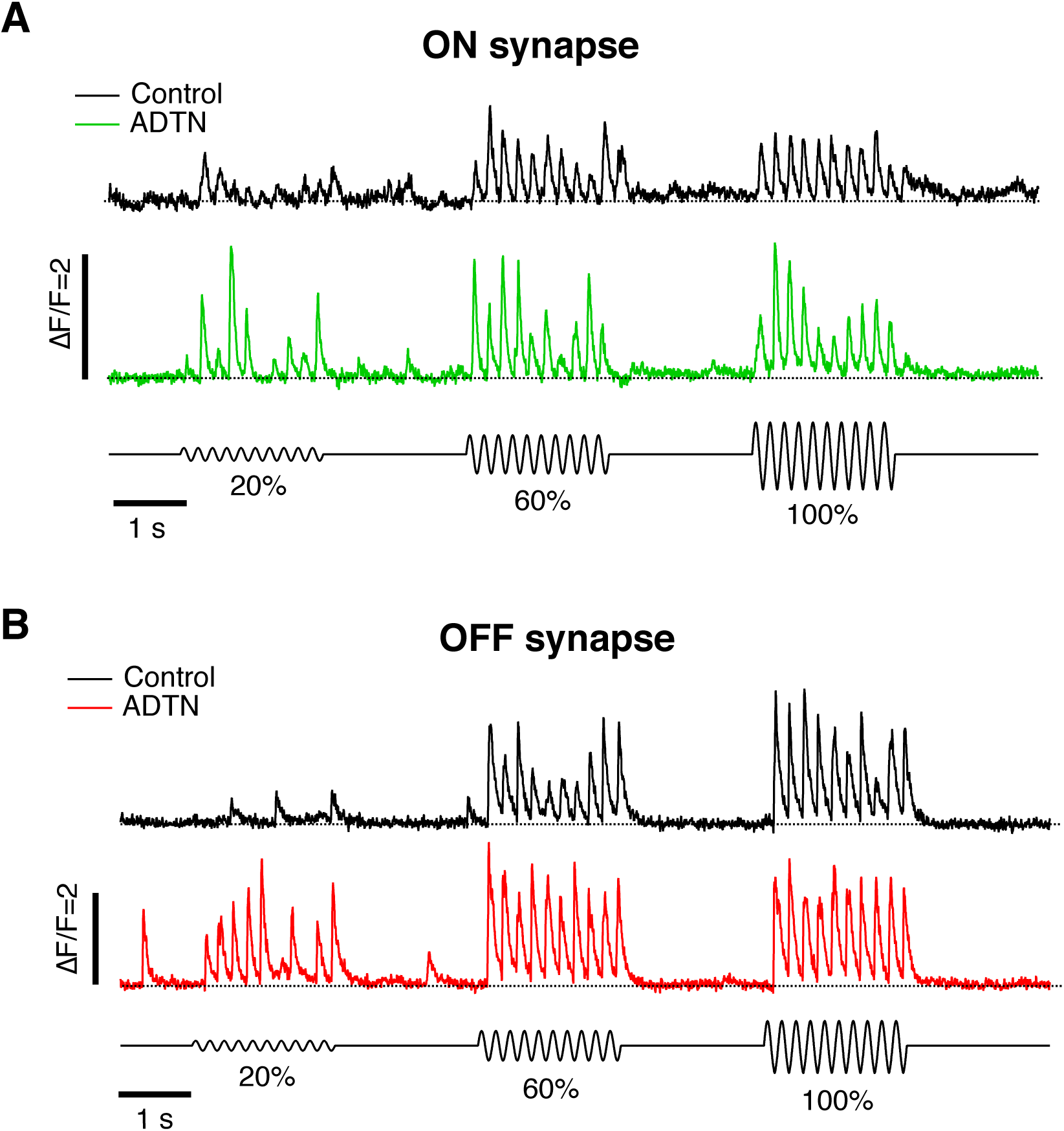
Activation of D1 receptors increased synaptic gain in both ON and OFF synapses. Examples of iGluSnFR signals from an ON synapse (A) and OFF synapse (B). The top trace (black) shows the response to contrasts of 20%, 60% and 100% and the coloured trace shows the responses to the same stimuli after injection of ADTN to an estimated final concentration of 200 nM. Similar observations were made in 15 OFF and 9 ON synapses.

**Figure S4.**
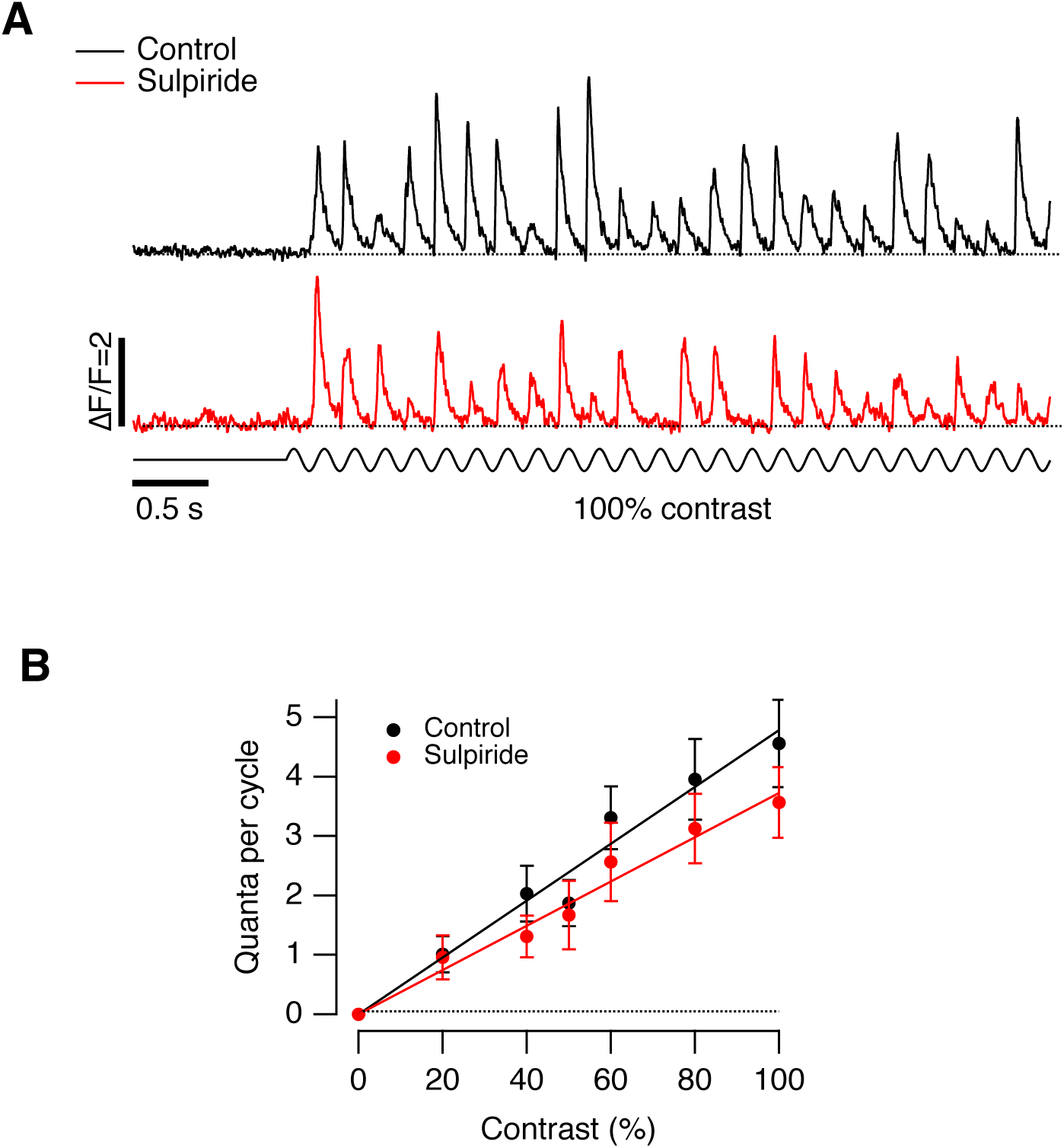
Antagonism of D2 receptors had a weak effect on synaptic gain. **A.** Examples of iGluSnFR signals from a synapse stimulated at 100% contrast before (black) and after (red) intravitreal injection of the D2 receptor antagonist Sulpiride to an estimated final concentration of 400 nM. **B.** Average contrast-response function from 10 paired recordings before and after intravitreal injection of Sulpiride. The measurements are described by a line through the origin. In control, the slope was 0.0488 ± 0.0005 and after injection of Sulpiride it was 0.037 ± 0.001, a difference significant at p < 0.0001 (t-test). From these slopes, the average reduction in synaptic gain was 28%.

**Figure S5.**
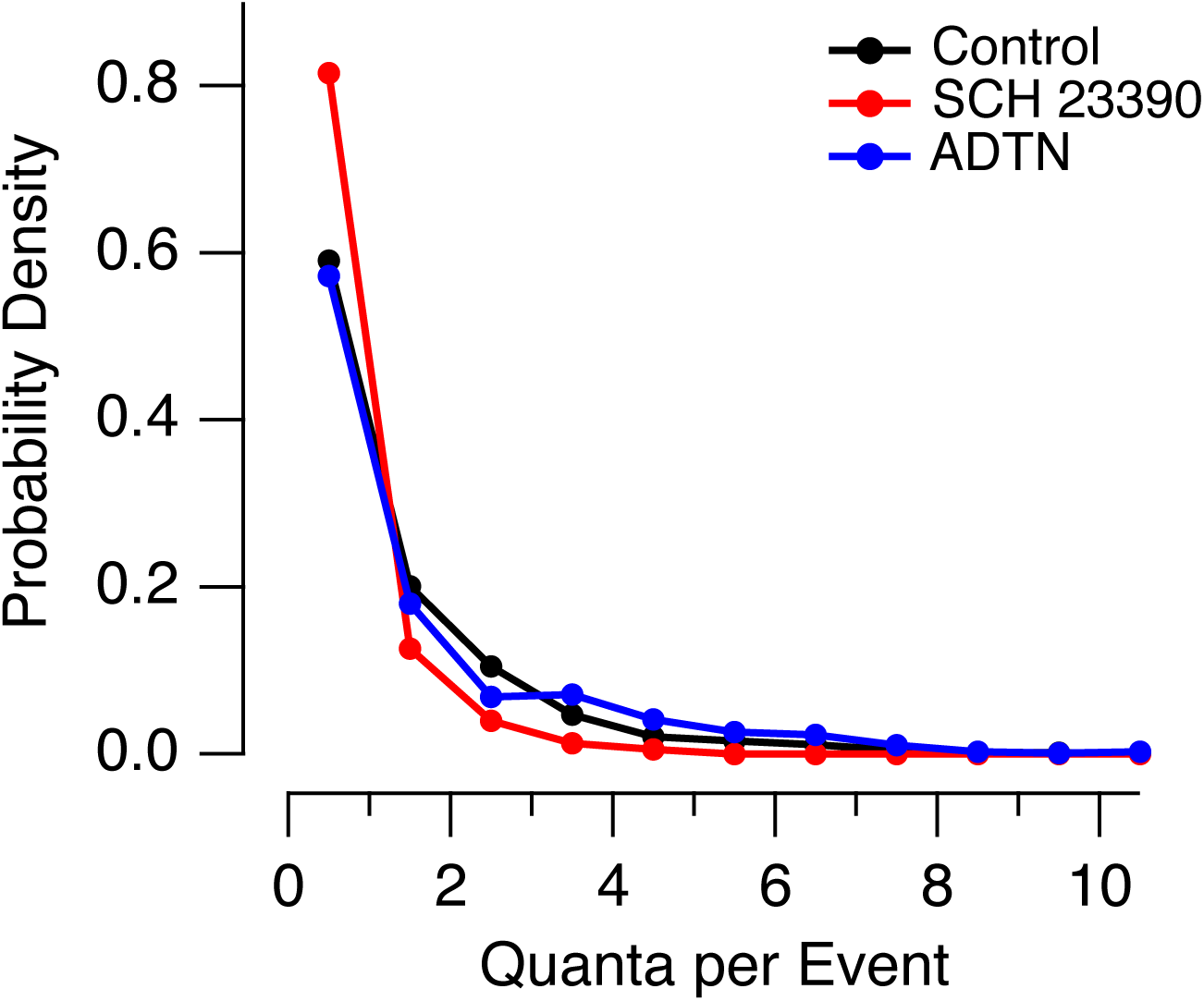
The effects of dopamine signaling on the amplitude of events triggered at low contrast. Changes in the distribution of Q_e_ (quanta per event) elicited at 20% contrast in control (n = 19 synapses), SCH 23390 (n = 10) and ADTN (n = 8). 5 Hz stimulus. The distribution in SCH 23390 was different to that observed under control conditions (p<0.001, Chi-squared test).

**Figure S6.**
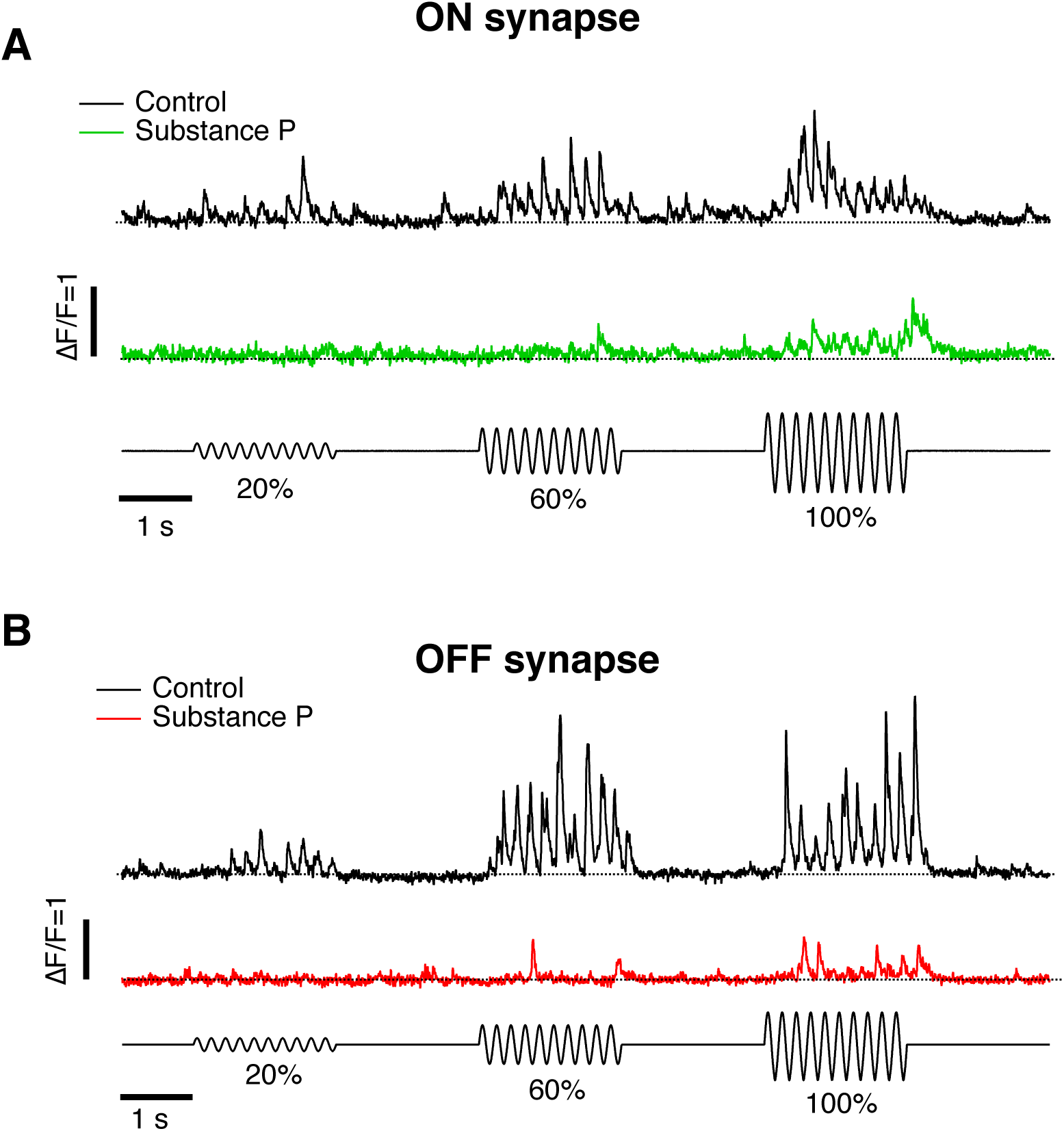
Activation of NK-1 receptors decreased synaptic gain in both ON and OFF synapses. Examples of iGluSnFR signals from an ON synapse (A) and OFF synapse (B). The top trace (black) shows the response to contrasts of 20%, 60% and 100% and the coloured trace shows the responses to the same stimuli after injection of Substance P to an estimated final concentration of 200 nM. Similar observations were made in 17 OFF and 7 ON synapses.

